# Differences in environmental stability among SARS-CoV-2 variants of concern: Omicron has higher stability

**DOI:** 10.1101/2022.01.18.476607

**Authors:** Ryohei Hirose, Yoshito Itoh, Hiroshi Ikegaya, Hajime Miyazaki, Naoto Watanabe, Takuma Yoshida, Risa Bandou, Tomo Daidoji, Takaaki Nakaya

## Abstract

SARS-CoV-2 variants of concern (VOCs) could cause significant human and economic damage owing to increased infectivity and transmissibility, and understanding their characteristics is crucial for infection control. Here, we analyzed differences in viral stability and disinfection efficacy between the Wuhan strain and all VOCs. On plastic and skin surfaces, Alpha, Beta, Delta, and Omicron variants exhibited more than two-fold longer survival than the Wuhan strain, and the Omicron variant had the longest survival time. Specifically, survival times of the Wuhan strain, Alpha variant, Beta variant, Gamma variant, Delta variant, and Omicron variant on skin surfaces were 8.6 h (95% CI, 6.5–10.9 h), 19.6 h (95% CI, 14.8–25.3 h), 19.1 h (95% CI, 13.9– 25.3 h), 11.0 h (95% CI, 8.1–14.7 h), 16.8 h (95% CI, 13.1–21.1 h), and 21.1 h (95% CI, 15.8– 27.6 h), respectively. *In vitro*, disinfectant effectiveness evaluations showed that Alpha, Beta, Delta, and Omicron were slightly more resistant to ethanol than the Wuhan strain. However, *ex vivo* evaluation showed that on human skin, all viruses were completely inactivated by exposure to 35 w/w % ethanol for 15 s. The high environmental stability of these VOCs could increase transmission risk and contribute to spread. Additionally, the Omicron variant might have been replaced by the Delta variant due to its increased environmental stability and rapid spread. To prevent VOC spread, it is highly recommended that current infection control practices use disinfectants with appropriate ethanol concentrations.

## Introduction

Various SARS-CoV-2 variants have emerged from 2020 to 2021. In particular, SARS-CoV-2 variants classified as variants of concern (VOCs) can cause significant human and economic damage, and an understanding their characteristics is crucial for infection control. VOCs have been reported to have increased infectivity and transmissibility (1). In particular, the rapid spread of the Omicron (Pango lineage: B.1.1.529) variant has become a serious concern worldwide as of 2022 (2, 3). The increase in the infectivity/transmissibility can be attributed to several factors, such as increased viral load shed from infected individuals, prolonged viral shedding period, decrease in the minimum viral load required to establish infection, changes in infection target site, and increased environmental stability (4, 5).

The environmental stability of SARS-CoV-2 has been compared with that of other viruses, such as SARS-CoV-1 and influenza virus (6, 7). Moreover, previous studies have suggested that Alpha (Pango lineage: B.1.1.7) and Beta (Pango lineage: B.1.351) variants have the same degree of environmental stability (8, 9). However, the differences in viral stability among all VOCs, including the Omicron and Delta (Pango lineage: B.1.617.2) variant, have not been evaluated and compared in detail. Here, we improved our previously developed evaluation model and precisely analyzed the differences in viral stability and disinfection efficacy between the Wuhan strain (Pango lineage: A) and all VOCs.

## Materials and methods

### Viruses and cells

The SARS-CoV-2 variants analyzed in this study were the Wuhan strain (Pango lineage: A, hCoV-19/Japan/TY/WK-521/2019), Alpha variant (Pango lineage: B.1.1.7, hCoV-19/Japan/QK002/2020), Beta variant (Pango lineage: B.1.351, hCoV-19/Japan/TY8-612/2021), Gamma variant (Pango lineage: P.1, hCoV-19/Japan/TY7-501/2021), Delta variant (Pango lineage: B.1.617.2, hCoV-19/Japan/TY11-927/2021), and Omicron variant (Pango lineage: B.1.1.529, hCoV-19/Japan/TY38-873/2021). All viruses were generously provided by the National Institute of Infectious Diseases (Tokyo, Japan). For virus culture and quantification, VeroE6/TMPRSS2 cells, expressing the transmembrane serine protease TMPRSS2, were purchased from the Japanese Collection of Research Bioresources Cell Bank (Osaka, Japan) and cultured in Dulbecco’s modified Eagle’s medium (DMEM; Sigma Aldrich) supplemented with 5% fetal bovine serum and G418 (Nacalai Tesque, Kyoto, Japan) (10, 11). The viruses were concentrated and purified as follows: 96 h post-infection, the culture medium was harvested and centrifuged for 10 min at 2,500 × *g* at 4 °C to eliminate cellular debris. Virions in the supernatant were sedimented through a 20% (w/w) sucrose cushion in phosphate-buffered saline (PBS) via ultracentrifugation at 27,000 rpm for 2.5 h at 4 °C using a Beckman SW28 rotor (12, 13). The titers of the virus were measured in terms of 50% tissue culture infectious dose (TCID_50_) in VeroE6/TMPRSS2 cells. Three days after inoculation, the cytopathic effect in each well was scored under a microscope, and the TCID_50_ was calculated.

### Construction of skin model to evaluate virus stability and disinfectant effectiveness

Human skin was collected from forensic autopsy specimens obtained from the Department of Forensic Medicine, Kyoto Prefectural University of Medicine. Abdominal skin autopsy specimens from subjects aged 20 to 70 years, obtained approximately 1 d after death, were cut into squares with approximate dimensions of 4 cm × 8 cm. Those whose skin was considerably damaged by burning or drowning were excluded. Using the skin-autopsy specimens, an *ex vivo* model was developed to evaluate the stability of different viruses on the surface of human skin and the effectiveness of different disinfectants against viruses on human skin (6, 14). The skin from which the panniculus adiposus that had been removed was washed with PBS and placed in a culture insert (Corning, Corning, NY, USA) on a membrane with a pore size of 8.0 μm. The culture inserts were placed in six-well plates containing 1.0 mL of DMEM (Sigma-Aldrich).

### Evaluation of virus stability on plastic and human skin surfaces

Virus survival was evaluated on plastic (polystyrene plate) and human skin (constructed skin model) surfaces. Virus solutions (5.0 × 10^4^ TCID_50_ in 2 μL PBS) were applied on the surface of plastic or human skin. Each sample was incubated in a controlled environment (25 °C, 45–55% relative humidity) for 0–120 h. The virus remaining on the surface was then collected in 1.0 mL of DMEM and titrated (6). The detection limit for the titer of the virus remaining on the surface was 10^0.5^ TCID_50_. Survival time was defined as the time until the virus on the surface was no longer detected. Three independent experiments were performed for each condition, and the results of residual virus titers on the surfaces were expressed as the mean ± standard error of the mean.

### Evaluation of effectiveness of alcohol-based disinfectants

The effectiveness of the ethanol-based disinfectants was evaluated at different concentrations. The effectiveness of ethanol (EA, Nacalai Tesque) was tested at concentrations of 80%, 60%, 50%, 40%, 35%, 32.5%, 30%, 27.5%, 25%, 22.5% and 20% (w/w). Isopropanol (IPA, Nacalai Tesque) was tested at a concentration of 70% (w/w).

First, an *in vitro* evaluation of the effectiveness of disinfectant was performed. In a 1500 μL tube; 5 μL of PBS containing virus (5.0 × 10^4^ TCID_50_ in 5 μL PBS) was mixed with 45 μL of various disinfectants for 15 s. Subsequently, the resulting solutions were neutralized with 450 μL of DMEM, and the remaining viral titers were measured (14). The detection limit for the virus titers was 10^0.2^ TCID_50_.

Next, the effectiveness of disinfectants against viruses on human skin was evaluated using the constructed model *(ex vivo* evaluation). Each virus solution (1.0 × 10^5^ TCID_50_ in 2 μL PBS) was applied to the human skin surface. Each skin sample was then incubated for 15 min at 25 °C with 45%–55% relative humidity to completely dry the viral mixture on the skin. Subsequently, 18 μL of disinfectant was applied to each skin sample surface for 15 s and then air-dried for 5 min. After drying, the remaining viruses on the skin were recovered with 1000 μL of DMEM, and the remaining viral load was measured (14). The detection limit for the virus titers was 10^0.5^ TCID_50_. To determine the effectiveness of the disinfectants under each condition, logarithmic reductions in virus titers were calculated, normalizing to the PBS control titers. Three independent experiments were performed for each condition, and the results were expressed as the mean ± standard error of the mean.

### Ethical considerations

The study protocol, including sample collection procedures, was reviewed and approved by the Institutional Review Board of the Kyoto Prefectural University of Medicine (ERB-C-1593). Written informed consent was obtained from all the study participants.

### Statistical analysis

Data were analyzed using GraphPad Prism 7 (GraphPad, Inc., La Jolla, CA, USA). The elapsed time was defined as an explanatory variable (X-axis), and the log virus titer was defined as an explained variable (Y-axis). Linear regression analysis with a logarithmic link function was performed to create a regression curve. The measurement limit of the SARS-CoV-2 titer was 10^0.5^ TCID_50_; therefore, the survival time was defined as the X-value when the Y-values of the regression curves were 0.5. The half-life was calculated from the slope of each regression curve when the titers of virus remaining on the surface were 2.0 and 3.0 log10TCID_50_ (6, 12).

## Results

The virus titers remaining on the plastic or human skin surfaces were measured over time (Supplementary Figure S1), and the survival time and half-life were calculated from these titer values by regression analysis (Supplementary Figure S2). In the plastic surface analysis, survival times of the Wuhan strain, Alpha variant, Beta variant, Gamma variant, Delta variant, and Omicron variant were 56.0 h (95% confidence interval [CI], 39.0–76.7 h), 191.3 h (95% CI, 152.5–232.1 h), 156.6 h (95% CI, 122.7–192.9 h), 59.3 h (95% CI, 43.9–77.7 h), 114.0 h (95% CI, 91.3–139.1 h), and 193.5 h (95% CI, 153.1–236.2 h), respectively (Figure 1A and Table 1). In the human skin surface analysis, survival times of the Wuhan strain, Alpha variant, Beta variant, Gamma variant, Delta variant, and Omicron variant were 8.6 h (95% CI, 6.5–10.9 h), 19.6 h (95% CI, 14.8–25.3 h), 19.1 h (95% CI, 13.9–25.3 h), 11.0 h (95% CI, 8.1–14.7 h), 16.8 h (95% CI, 13.1–21.1 h), and 21.1 h (95% CI, 15.8–27.6 h), respectively (Figure 1B and Table 1). Alpha, Beta, Delta, and Omicron variants had significantly longer survival times than the Wuhan strain, and the Omicron variant had the longest survival time. Furthermore, the half-life showed the same tendency as the survival time (Figure 1C and 1D and Table 2).

**Figure 1.**
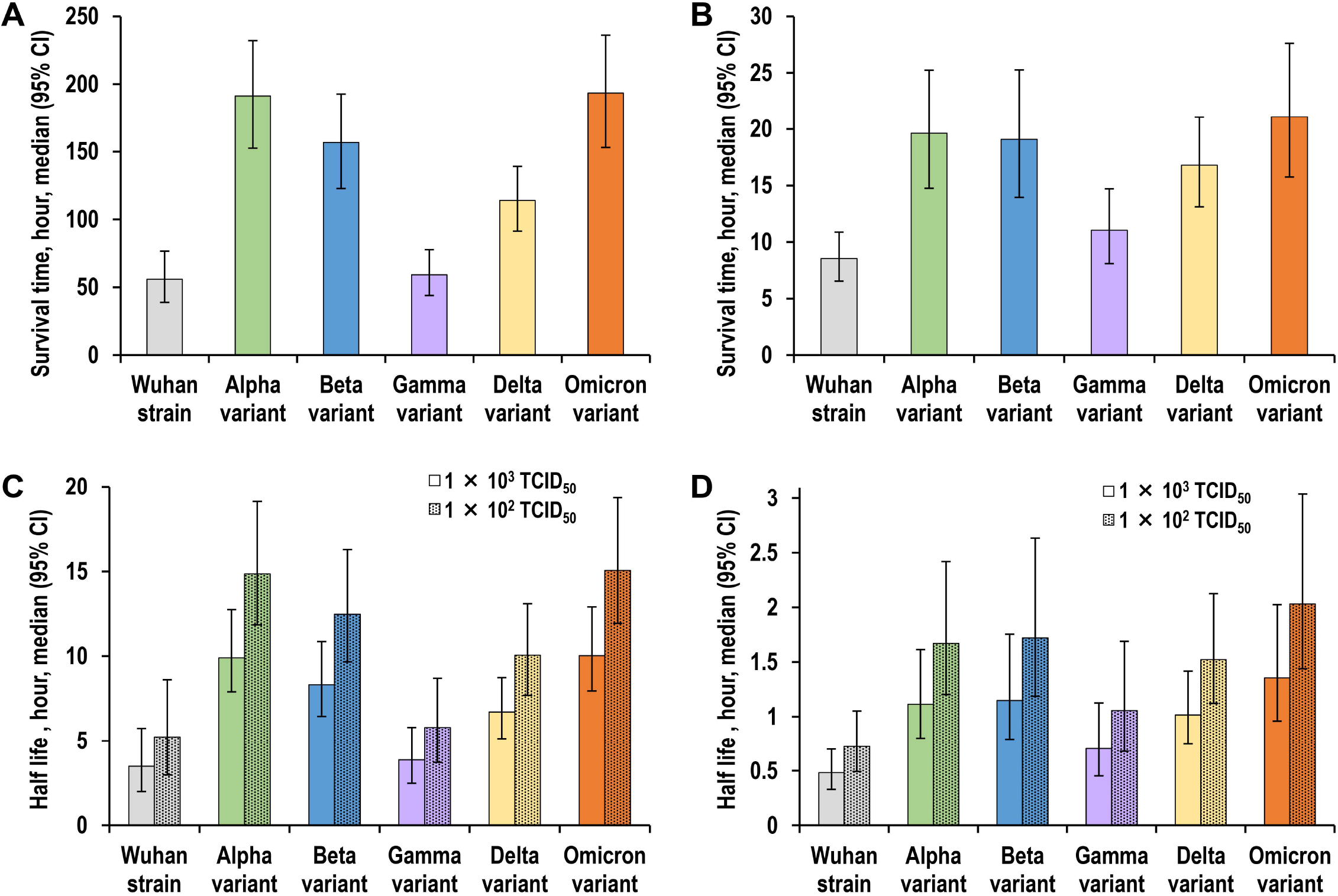
(A) Survival times of the various viruses on a plastic surface. (B) Survival times of the various viruses on the surface of human skin. (C) Half-lives of the various viruses on a plastic surface. (D) Half-lives of the various viruses on the surface of human skin. Survival time is defined as the time until the virus was no longer detected on the surface. All half-lives in the graphs refer to a condition when 1 × 10^2^ or 1 × 10^3^ TCID_50_ (50% tissue culture infectious dose) of virus particles remain on the surface. Data are expressed as the median ± 95% confidence interval.

**Table 1.** Survival time of SARS-CoV-2 on plastic and human skin surfaces.

|  | Survival time, hour, median (95% CI) |  |
| --- | --- | --- |
|  | Plastic surface | Skin surface |
| **Wuhan strain** | 56.0 (39.0-76.7) | 8.6 (6.5-10.9) |
| **Alpha variant** | 191.3 (152.5-232.1) | 19.6 (14.8-25.3) |
| **Beta variant** | 156.6 (122.7-192.9) | 19.1 (13.9-25.3) |
| **Gamma variant** | 59.3 (43.9-77.7) | 11.0 (8.1-14.7) |
| **Delta variant** | 114.0 (91.3-139.1) | 16.8 (13.1-21.1) |
| **Omicron variant** | 193.5 (153.1-236.2) | 21.1 (15.8-27.6) |

The elapsed time was defined as an explanatory variable (X-axis), and the log virus titer was defined as an explained variable (Y-axis). A linear regression analysis with logarithmic link function was performed to create a curve of regression. The measurement limit of the SARS-CoV-2 titer was $10^{0.5}$ TCID<sub>50</sub>; therefore, the survival time was defined as the X values when the Y values of the regression curves were 0.5.

**Table 2.** Half-life of SARS-CoV-2 on the plastic and human skin surfaces.

|  | Half-life, hour, median (95% CI) |  |  |  |
| --- | --- | --- | --- | --- |
|  | Plastic surface |  | Skin surface |  |
|  | 3 (Log<sub>10</sub>TCID<sub>50</sub>) | 2 (Log<sub>10</sub>TCID<sub>50</sub>) | 3 (Log<sub>10</sub>TCID<sub>50</sub>) | 2 (Log<sub>10</sub>TCID<sub>50</sub>) |
| **Wuhan strain** | 3.5 (2.0-5.7) | 5.2 (3.0-8.6) | 0.5 (0.3-0.7) | 0.7 (0.5-1.1) |
| **Alpha variant** | 9.9 (7.9-12.7) | 14.9 (11.9-19.1) | 1.1 (0.8-1.6) | 1.7 (1.2-2.4) |
| **Beta variant** | 8.3 (6.4-10.9) | 12.5 (9.7-16.3) | 1.2 (0.8-1.8) | 1.7 (1.2-2.6) |
| **Gamma variant** | 3.9 (2.5-5.8) | 5.8 (3.7-8.7) | 0.7 (0.5-1.1) | 1.1 (0.7-1.7) |
| **Delta variant** | 6.7 (5.1-8.7) | 10.1 (7.7-13.1) | 1.0 (0.8-1.4) | 1.5 (1.1-2.1) |
| **Omicron variant** | 10.0 (8.0-12.9) | 15.1 (11.9-19.4) | 1.4 (1.0-2.0) | 2.0 (1.4-3.0) |

The elapsed time was defined as an explanatory variable (X-axis), and the log virus titer was defined as an explained variable (Y-axis). A linear regression analysis with logarithmic link function was performed to create a curve of regression. The half-life was calculated from the slope of each regression curve when the titer of virus remaining on the surface was 2.0, and 3.0 Log<sub>10</sub>TCID<sub>50</sub>.

The *in vitro* disinfectant effectiveness evaluation showed that the Wuhan strain and Gamma variant were completely inactivated within 15 s by 32.5% EA (log reduction > 4), Alpha, Beta, and Delta variants were completely inactivated within 15 s by 35% EA, and the Omicron variant was completely inactivated within 15 s by 40% EA (Figure 2A and Supplementary Table S1). Alpha, Beta, Delta, and Omicron variants were thus slightly more resistant to ethanol than the Wuhan strain. However, on human skin, an *ex vivo* evaluation showed that all viruses were completely inactivated after exposure to 35% EA for 15 s (log reduction > 4; Figure 2B and Supplementary Table S2).

**Figure 2.**
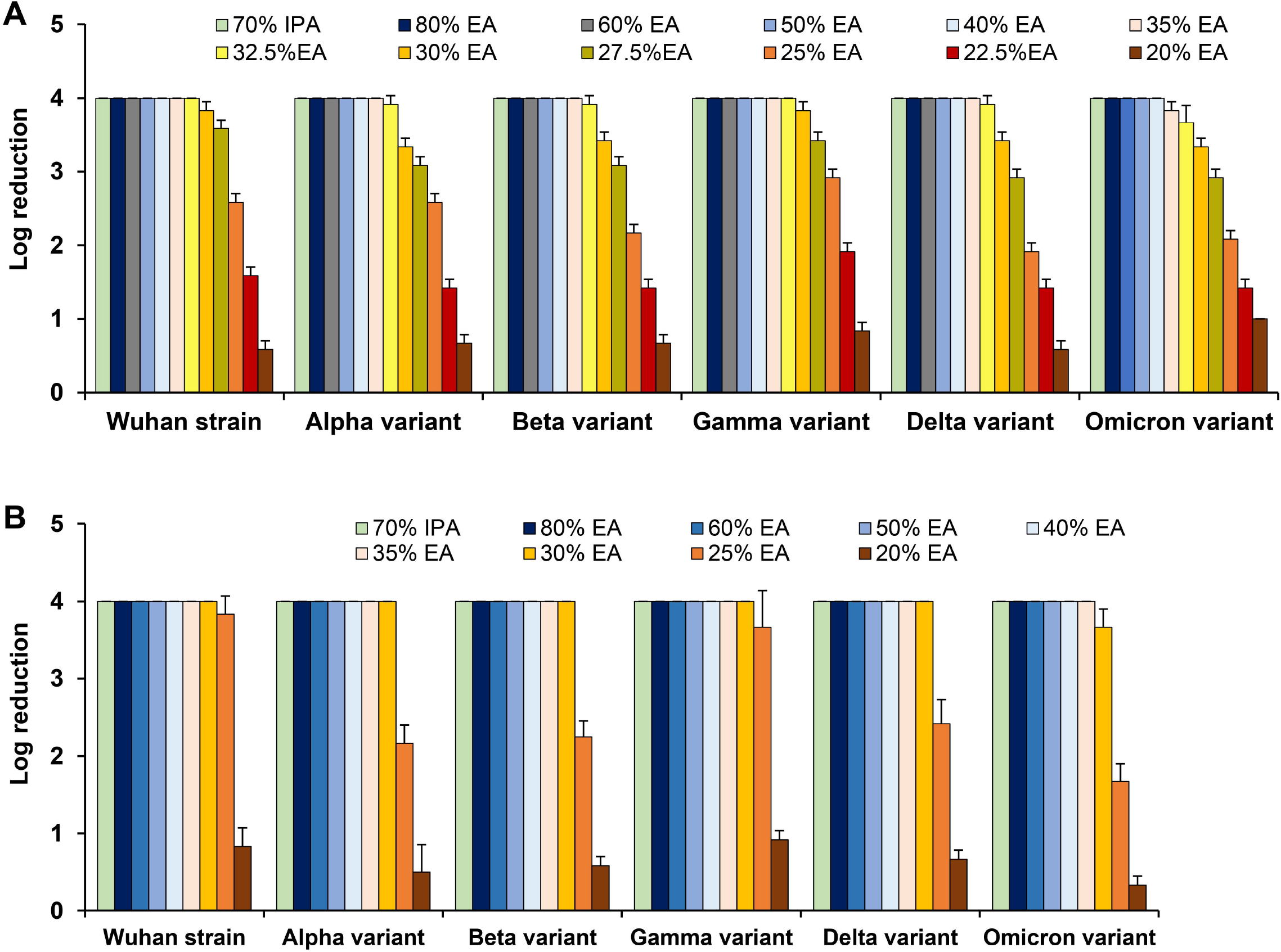
Evaluation of disinfectant effectiveness. Both *in vitro* evaluation (A) and *ex vivo* evaluation (B) were performed, and the log reduction was calculated from the residual viral titer after each alcohol-based disinfectant exposure (see Supplementary Table S1 and S2). The results are expressed as the mean ± standard error. EA, ethyl alcohol; IPA, isopropanol.

## Discussion

In 2020, the environmental stability of the Wuhan strain was reported in several studies (6, 7, 15). Additionally, some studies suggested that the Alpha and Beta variants have the same degree of environmental stability (8, 9). However, no study has directly compared other VOCs with the Wuhan strain, and the differences in environmental stability between the Wuhan strain and all VOCs, including Omicron and Delta variants, were previously unknown.

Our study showed that on plastic and skin surfaces, Alpha, Beta, Delta, and Omicron variants exhibited more than two-fold longer survival times than those of the Wuhan strain and maintained infectivity for more than 16 h on the skin surfaces. The high environmental stability of these VOCs could increase the risk of contact transmission and contribute to the spread of VOCs. Additionally, in this analysis, there was no significant difference in survival times between Alpha and Beta variants, and they had similar environmental stability, which is consistent with the results of previous studies (8, 9).

The Omicron variant is currently a major concern owing to the rapidly increasing number of infected patients worldwide. The shift in the target site of infection from the lower respiratory tract to the upper respiratory tract and the escape from neutralizing antibodies might be potential factors for the spread of the Omicron variant (1–5). This study showed that the Omicron variant also has the highest environmental stability among VOCs, which suggests that this high stability might also be one of the factors that have allowed the Omicron variant to replace the Delta variant and spread rapidly. Although Alpha, Beta, Delta, and Omicron variants showed a slight increase in ethanol resistance in response to increased environmental stability, all VOCs on the skin surface were completely inactivated by 15 s exposure to 35% EA. Therefore, it is highly recommended that current infection control (hand hygiene) practices use disinfectants with appropriate EA concentrations (>52 w/w% or >60 v/v%), as proposed by the World Health Organization (16, 17).

This study had three limitations. First, the reason for the higher environmental stability of Alpha, Beta, Delta, and Omicron variants is unknown at this stage, and evaluation using recombinant viruses might identify factors that determine this. Second, the survival time and half-life obtained in this environmental stability evaluation might vary depending on the external environment and the composition of the body fluid containing the virus. In this study, to accurately analyze the differences in stability between VOCs, the target virus was purified by ultracentrifugation, and PBS was used as a solvent. Third, the relationship between the amount of virus remaining on the surface and the risk of transmission is still unclear at this stage. Therefore, it might be reasonable to interpret the value of survival time in this study as a reference value.

In conclusion, we elucidated the environmental stability of VOCs, which is important information for infection control. Furthermore, these findings will contribute greatly to elucidating the mechanism of VOC spread with the addition of genetic analysis.

## Acknowledgments

We thank Editage (www.editage.com) for English language editing. This research was supported by Adaptable and Seamless Technology Transfer Program through Target-driven R&D (ASTEP) from the Japan Science and Technology Agency (JST) [grant number JPMJTR21UE and JPMJTM20PR], JSPS KAKENHI (grant number 21K16326), Mitsubishi Foundation, and Takeda Science Foundation.

## Author contributions

Study concept and design: RH. Data acquisition: RH, HM, NW, TY, RB, TD. Data analysis and interpretation: RH, YI, and TN. Drafting of the manuscript: RH. Statistical analysis: RH. Secured funding: RH. Administrative/technical/material support: RH, HI. Study supervision: RH and TN.

