## Supplementary Material (Supplementary figures and tables)) for "Differences in environmental stability among SARS-CoV-2 variants of concern: Omicron has higher stability"

**This PDF file includes:**

1. Supplementary Figure S1.

2. Supplementary Figure S2.

3. Supplementary Table S1.

4. Supplementary Table S2.

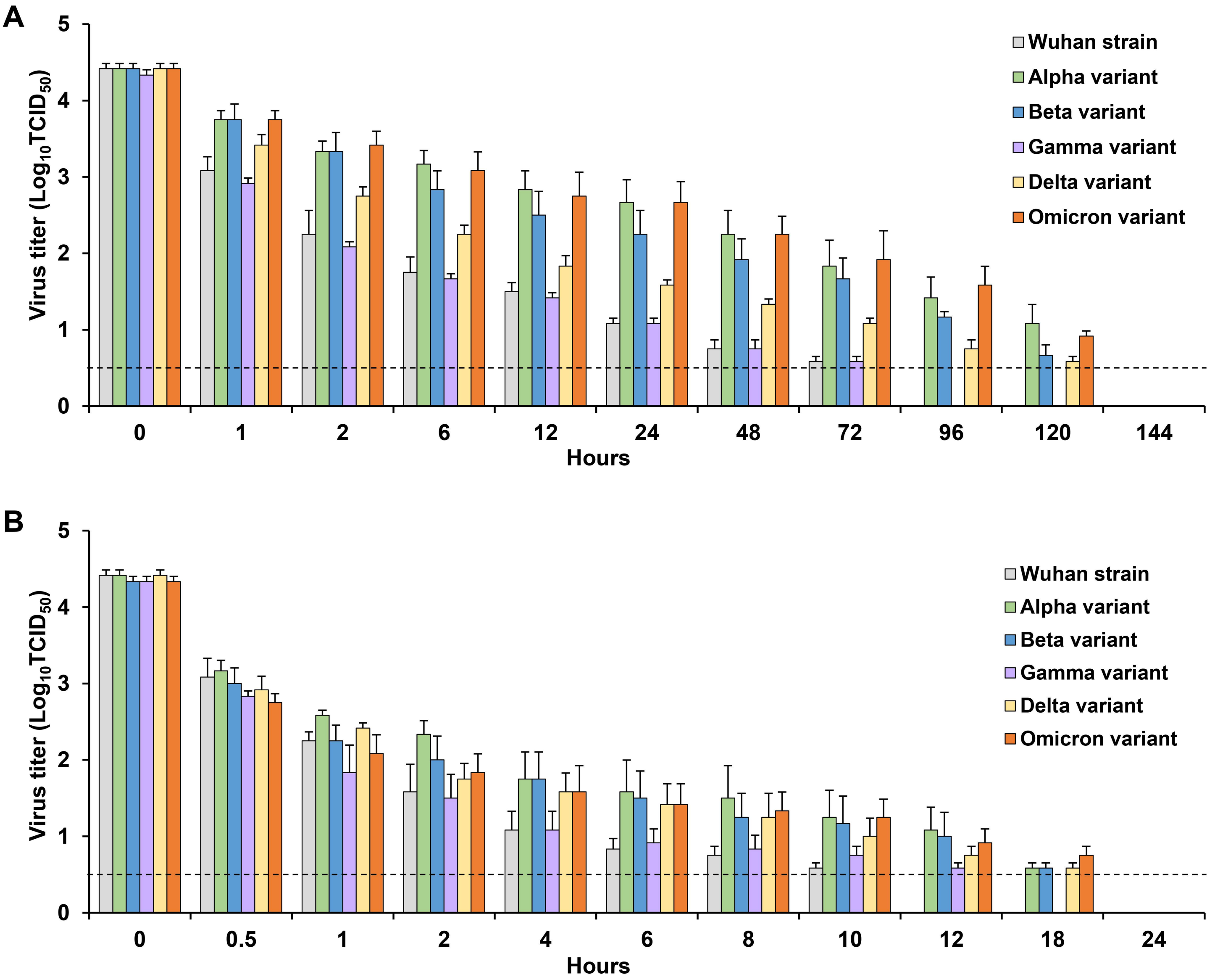


**Supplementary Figure S1. Decrease in the residual viral titer on the plastic surface (A) and human skin surface (B) as a function of time.** Each virus ($5.0 \times 10^4$ $TCID_{50}$; 50% tissue culture infectious dose) was mixed with 2 μL of phosphate-buffered saline and applied to each surface. Each surface was incubated in a controlled environment (temperature: 25 °C, humidity: 45%–55%) for 0–144 h. The virus on the surface was then recovered in 1 mL of medium and titrated to calculate the titer of the virus remaining on the surface. For each condition, three independent experiments were performed, and the results are expressed as the mean ± standard error of the mean. The bars for data below the detection limit were omitted. The dotted horizontal lines represent the detection limit titers.

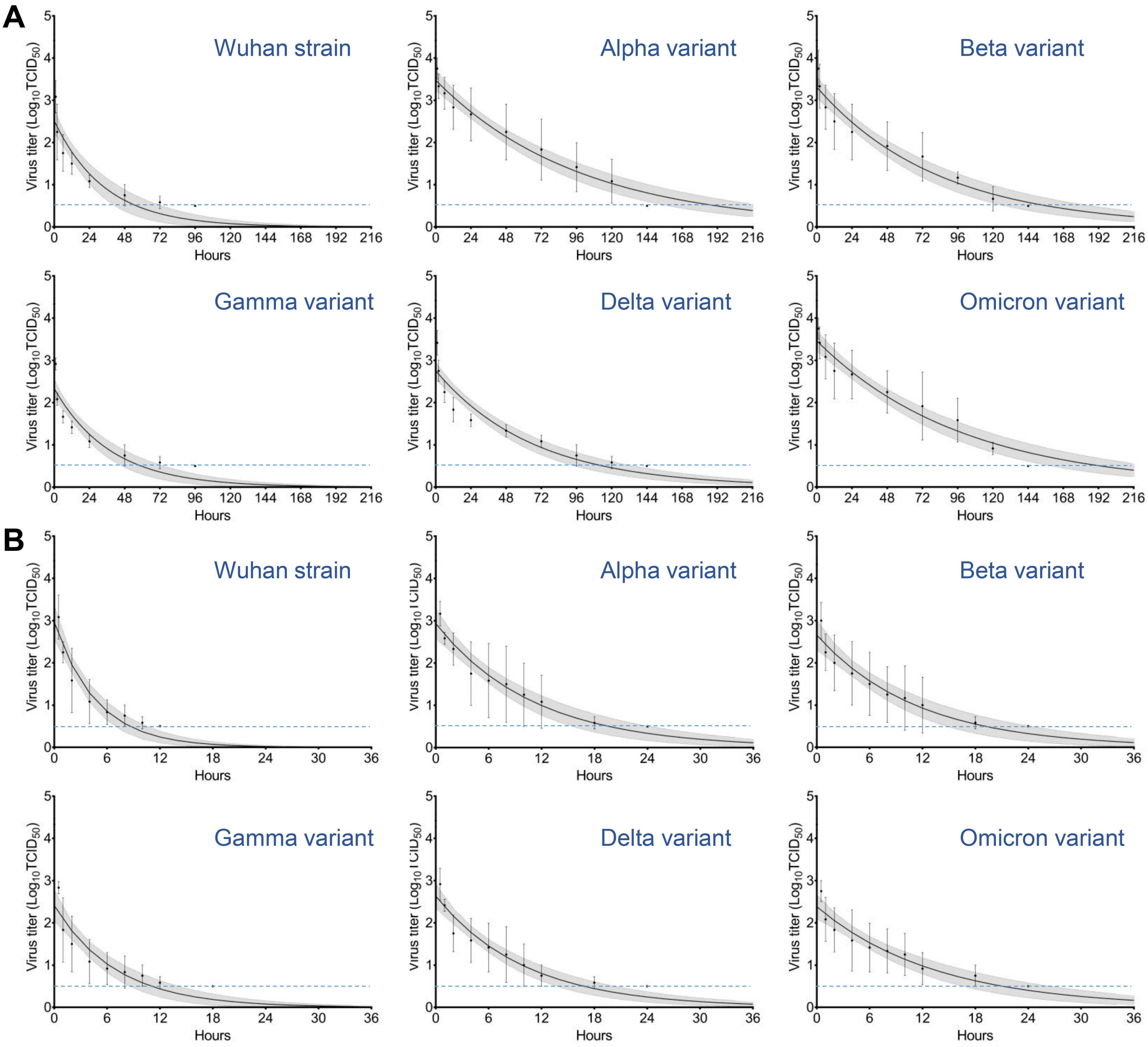


**Supplementary Figure S2. Stability of viruses on the plastic surface (A) and the human skin surface (B).** The elapsed time was defined as an explanatory variable (X-axis), and the log virus titer was defined as an explained variable (Y-axis). Least-squares linear-regression analysis was performed with a logarithmic link function for each virus to generate a regression curve. The upper and lower confidence limits are represented by dotted curves. The dotted horizontal lines represent the detection limit titers. The data shown are expressed as the mean ± standard error of the mean for three independent experiments.

**Supplementary Table S1.** ***In vitro*** **evaluation of disinfectant effectiveness.**

| | Log reduction, mean ± standard error | | | | | |
|---|---|---|---|---|---|---|
| | **Wuhan strain** | **Alpha variant** | **Beta variant** | **Gamm variant** | **Delta variant** | **Omicron variant** |
| **70% IPA** | 4.00 | 4.00 | 4.00 | 4.00 | 4.00 | 4.00 |
| **80% EA** | 4.00 | 4.00 | 4.00 | 4.00 | 4.00 | 4.00 |
| **60% EA** | 4.00 | 4.00 | 4.00 | 4.00 | 4.00 | 4.00 |
| **50% EA** | 4.00 | 4.00 | 4.00 | 4.00 | 4.00 | 4.00 |
| **40% EA** | 4.00 | 4.00 | 4.00 | 4.00 | 4.00 | 4.00 |
| **35% EA** | 4.00 | 4.00 | 4.00 | 4.00 | 4.00 | 3.83 ± 0.12 |
| **32.5%EA** | 4.00 | 3.92 ± 0.12 | 3.92 ± 0.12 | 4.00 | 3.92 ± 0.12 | 3.67 ± 0.24 |
| **30% EA** | 3.83 ± 0.12 | 3.33 ± 0.12 | 3.42 ± 0.12 | 3.83 ± 0.12 | 3.42 ± 0.12 | 3.33 ± 0.12 |
| **27.5%EA** | 3.58 ± 0.12 | 3.08 ± 0.12 | 3.08 ± 0.12 | 3.42 ± 0.12 | 2.92 ± 0.12 | 2.92 ± 0.12 |
| **25% EA** | 2.58 ± 0.12 | 2.58 ± 0.12 | 2.17 ± 0.12 | 2.92 ± 0.12 | 1.92 ± 0.12 | 2.08 ± 0.12 |
| **22.5%EA** | 1.58 ± 0.12 | 1.42 ± 0.12 | 1.42 ± 0.12 | 1.92 ± 0.12 | 1.42 ± 0.12 | 1.42 ± 0.12 |
| **20% EA** | 0.58 ± 0.12 | 0.67 ± 0.12 | 0.67 ± 0.12 | 0.83 ± 0.12 | 0.58 ± 0.12 | 1.00 ± 0.00 |

The log reduction value was calculated to evaluate disinfectant effectiveness under each condition and was expressed as mean ± standard error. EA, ethanol; IPA, isopropanol.

**Supplementary Table S2. Evaluation of disinfectant effectiveness against viruses on a human skin surface.**

| | Log reduction, mean ± standard error | | | | | |
|---|---|---|---|---|---|---|
| | **Wuhan strain** | **Alpha variant** | **Beta variant** | **Gamma variant** | **Delta variant** | **Omicron variant** |
| **70% IPA** | 4.00 | 4.00 | 4.00 | 4.00 | 4.00 | 4.00 |
| **80% EA** | 4.00 | 4.00 | 4.00 | 4.00 | 4.00 | 4.00 |
| **60% EA** | 4.00 | 4.00 | 4.00 | 4.00 | 4.00 | 4.00 |
| **50% EA** | 4.00 | 4.00 | 4.00 | 4.00 | 4.00 | 4.00 |
| **40% EA** | 4.00 | 4.00 | 4.00 | 4.00 | 4.00 | 4.00 |
| **35% EA** | 4.00 | 4.00 | 4.00 | 4.00 | 4.00 | 4.00 |
| **30% EA** | 4.00 | 4.00 | 4.00 | 4.00 | 4.00 | 3.67 ± 0.24 |
| **25% EA** | 3.83 ± 0.24 | 2.17 ± 0.24 | 2.25 ± 0.20 | 3.67 ± 0.47 | 2.42 ± 0.31 | 1.67 ± 0.24 |
| **20% EA** | 0.83 ± 0.24 | 0.50 ± 0.35 | 0.58 ± 0.12 | 0.92 ± 0.12 | 0.67 ± 0.12 | 0.33 ± 0.12 |

The log reduction value was calculated to evaluate disinfectant effectiveness under each condition and was expressed as mean ± standard error. EA, ethanol; IPA, isopropanol.
